# The modular biochemical reaction network structure of cellular translation

**DOI:** 10.1101/2023.01.21.524914

**Authors:** Bruno Cuevas-Zuviría, Evrim Fer, Zachary R. Adam, Betül Kaçar

## Abstract

Translation is an essential attribute of all living cells. At the heart of cellular operation, it is a chemical information decoding process that begins with an input string of nucleotides and ends with the synthesis of a specific output string of peptides. The translation process is interconnected with gene expression, physiological regulation, transcription, and responses to signaling molecules, among other cellular functions. Foundational efforts have uncovered a wealth of knowledge about the mechanistic functions of and many interactions between components of translation, but the broader biochemical connections between translation, metabolism and polymer biosynthesis that enable translation to occur have not been comprehensively mapped. Here we present a multilayer graph of biochemical reactions describing the translation, polymer biosynthesis and metabolism networks of an *Escherichia coli* cell. Intriguingly, the compounds that compose these three layers are distinctly aggregated into three modes regardless of their layer categorization. Multimodal mass distributions are well-known in ecosystems, but this is the first such distribution reported at the biochemical level. The degree distributions of the translation and metabolic networks are each likely to be heavy-tailed, but the polymer biosynthesis network is not. A multimodal mass-degree distribution indicates that the translation and metabolism networks are each distinct, adaptive biochemical modules, and that the gaps between the modes reflect evolved responses to the functional use of metabolite, polypeptide and polynucleotide compounds. The chemical reaction network of cellular translation opens new avenues for exploring complex adaptive phenomena such as percolation and phase changes in biochemical contexts.

## Introduction

Cellular life is a network of molecular components that are dynamic and adaptive. Life can only exist as a gradient of free energy, and metabolic networks can account for the uptake and excretion of substrates alongside the synthesis of key biomolecules^1^. Metabolic networks trace the routes that energy travels through cells, but genetic sequences and their associated enzymatic polymers^2–4^ specify which pathways will be constructed, governed, and maintained in any given cell. While a great deal of current research is focused on filling specific knowledge gaps at the biochemical level, studies of the overall cellular network can provide insights into how the details of biochemistry lead to the emergence of life’s foundational properties.

At the cellular level, analysis of metabolic networks has inspired decades of research into biochemical complexity^5–9^. These analyses have drawn connections between network attributes such as a heavy-tailed degree distributions and general complex behaviors such as resilience, adaptability, and modularity. Following initial studies of metabolic networks, there have been numerous descriptions of heavy-tailed networks at other biological levels such as ecological interactions, including theories of generative mechanisms that are relevant to characterizing the prebiotic world^10–14^. These generative mechanisms lead to continuous, heavy-tailed distributions consistent with a power law as an asymptotic sampling limit is approached; discontinuous or multimodal distributions are not explicitly prescribed by these physical models.

While the cellular level is usually the fundamental starting point to describe life properties, life manifests at different nested scales (*e.g.*, cell, organism, ecosystem). At the largest biological levels of populations and ecologies, systems show emergent traits as *multimodality* or even *discontinuities* in their properties. One of the most studied properties showing discontinuities is size: depending on the ecosystem, organisms spanning certain size ranges will not exist. Examples are found in arboreal forest^15^, bird^16^, fish and plankton^17^, and mammal^18^ groupings that can exhibit multimodal or discontinuous size distributions. Ecological multimodal aggregations arise from a confluence of dynamic processes consisting of an overlapping combination of both positive feedback (*e.g.*, overabundant energy resources available for consumption at one size or temporal scale but not at others) and negative feedback (*e.g.*, predation pressures such as grazing that can make survival at specific scales extremely challenging, or few and highly variable available resources)^19^. Discontinuities and modalities in biology also appear to be idiosyncratic; gaps or troughs in one ecosystem do not necessarily map to those found in others, and persistent gaps can nevertheless vary and shift over time^20^. It is not clear whether these network-level attributes of biology ought to extend downward to the biomolecular scale.

Considering these scale-dependent aspects of biological networks, a compelling possibility is which (if any) topological features associated with complex high-level biological systems can be found at the cellular level. We hypothesized that these features, if they show up within a cellular system, necessitate an explicit description of the processes that shape chemical hierarchical structure. In ecosystems, these processes include mapping trophic relationships and sources of energy input. Therefore, we address this question at the cellular level by building a translation reaction network in *Escherichia coli (E. coli)*, which we have combined with its metabolic and biosynthetic networks (**Figure 1**). Translation is a molecular information decoding process that begins with a ribonucleotide sequence as input and a peptide sequence as output, with remarkable robustness, accuracy and adaptability^21–27^. The processes that allow translation to occur are carried out through the coordinated actions of numerous ribonucleic and protein polymers, which we refer to as the translation machinery (TM). The decoding process that converts triplets of RNA monomers into the attachment of a specific amino acid residue to a growing peptide sequence is remarkable in that it is the earliest (and possibly only) emergent system that processes discretized information and manifests semantic relationships entirely in chemical terms. The chemosynthetic nature of the TM (*i.e.*, information processing via chemical reactions, as compared to electronic signals in the brain or in digital computers) enables its description as a chemical reaction network, which is amenable to systems-level analyses as has been done for metabolic networks^5, 28^.

**Figure 1.**
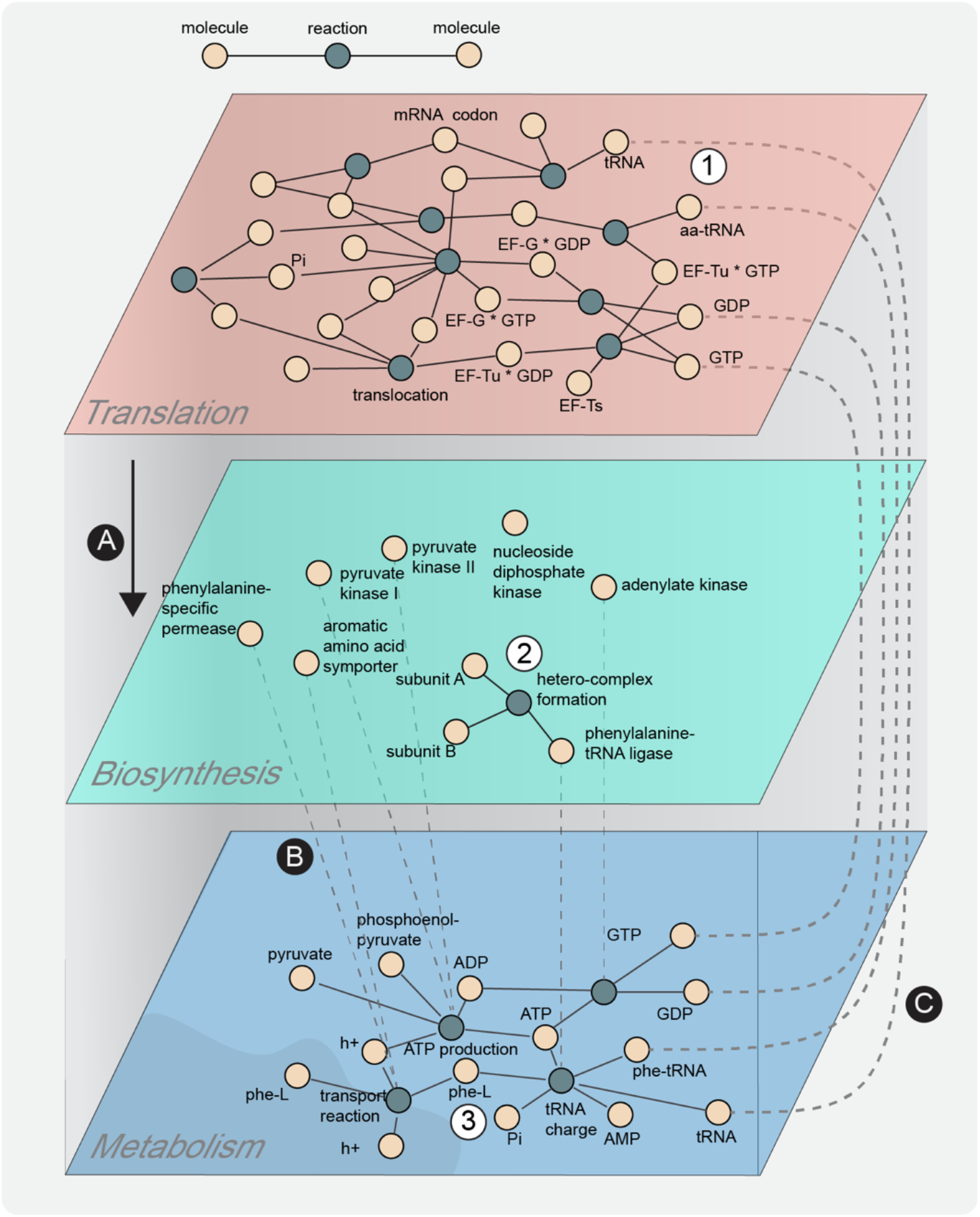
Schematic representation of the integrated cellular chemosynthesis network, composed of translation, polymer biosynthesis and metabolism layers. We built a translation layer (1) that includes all the relevant steps of that process such as initiation, peptidic bond formation, translocation, and termination. Translation generates peptides that appear in a biosynthesis layer (A). The biosynthesis layer describes interactions between polymers and the formation of reported hetero-complex multimers (2) to carry out metabolic functions. Proteins in the biosynthesis layer act as enzymes or transporters enabling reactions (B) that generate energy and building blocks of translation such as ATP, GTP and charged tRNA within the metabolism layer (3). Finally, the free energy and building blocks of metabolism are employed as substrate inputs for translation (C).

Here, we present the translation reaction network that includes the core processes of translation (initiation, elongation, termination), detailed assembly steps required to replicate the ribosome, and other well-documented features of the TM that maintain translation functionality (**Figure 2A**). To connect the translation and metabolic networks, we constructed a polymer biosynthesis network that explicitly links enzymatic components catalyzing the metabolic reactions with the mRNA reading and peptide synthesis steps of translation (**Figure 3A**). Comprehensive mapping of the connections between translation and metabolism is necessary to investigate context-dependent feedbacks between the cellular environment, substrate/nutrient availability, and ribosomal protein activity.

**Figure 2.**
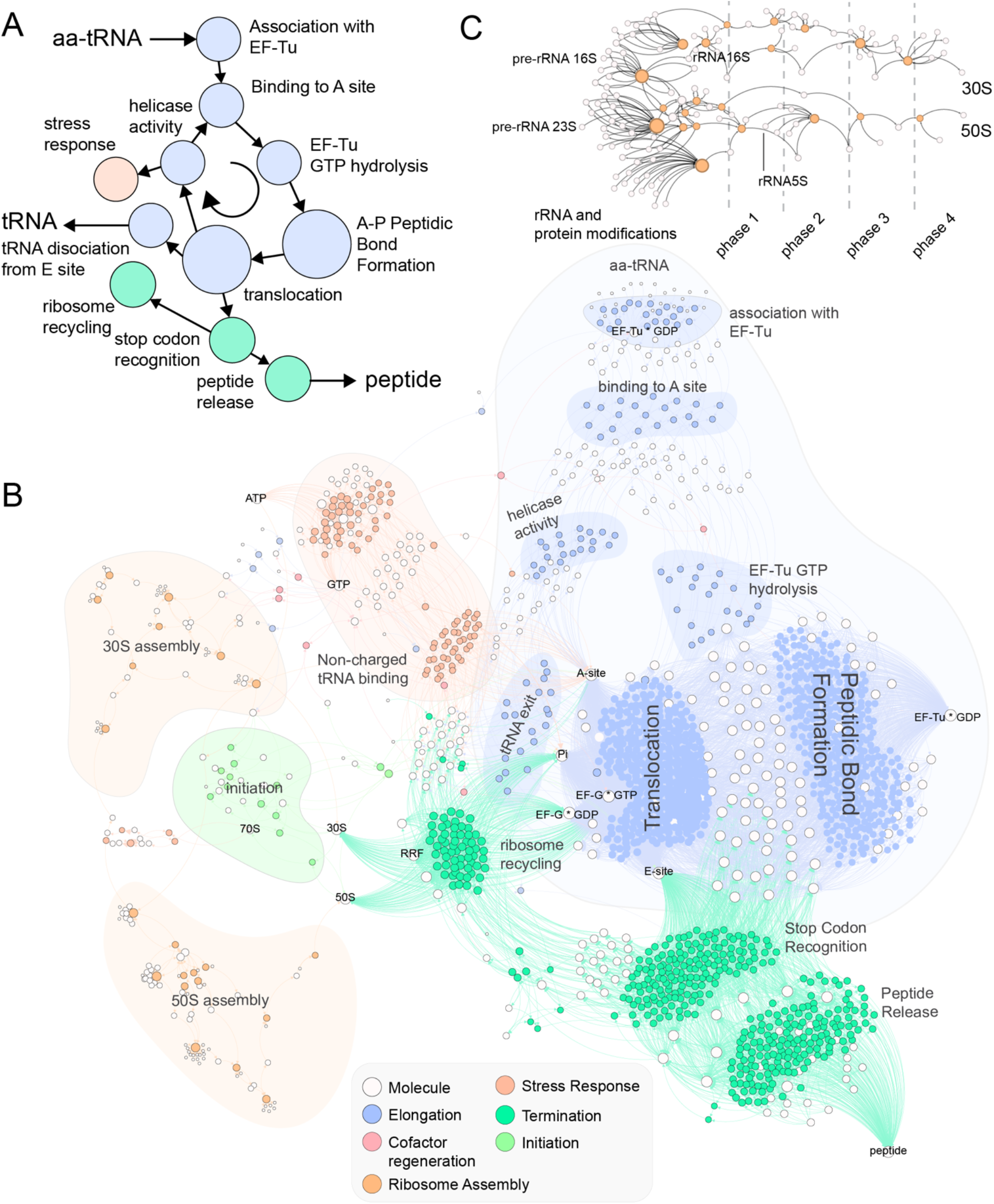
Overview of the translation portions of the integrated chemosynthesis network. . **A.** A simplified depiction of the ‘canonical’ translation process encoded within the translation network. **B.** A detailed visualization of the complete translation chemical reaction layer using Gephi 0.9.2 and a Force Atlas 2 layout^88^ (settings: scaling=5.0, gravity=1.0, tolerance=1.0, approximation=1.2). **C.** A depiction of the 30S and 50S component assembly steps that enable ribosomal replication to occur.

**Figure 3.**
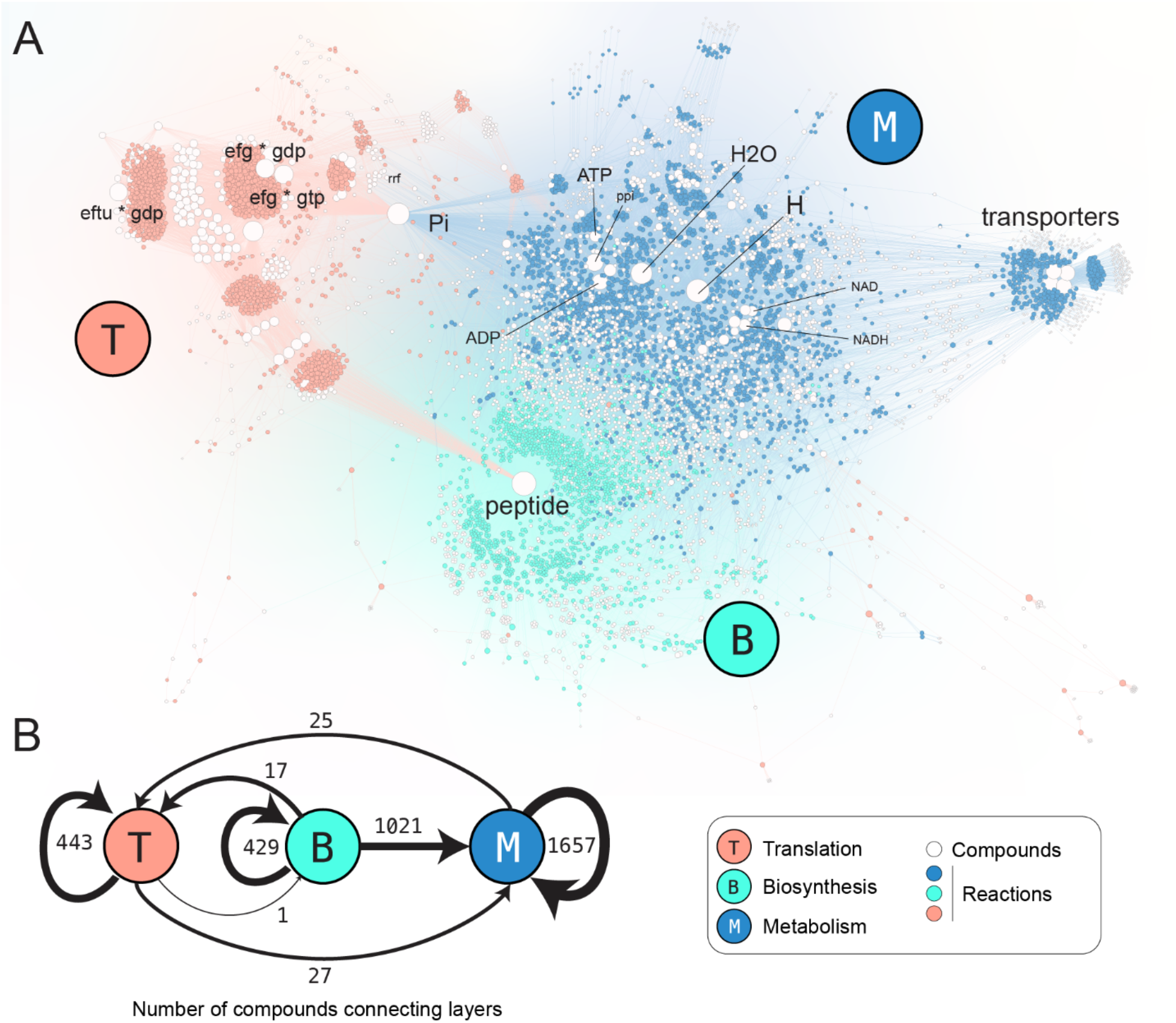
Detailed and simplified visualizations of all reactions and compounds that compose the integrated chemosynthesis network. **A.** Full map with Translation (T) reactions in orange, Polymer Biosynthesis (B) reactions in green, and Metabolic (M) reactions in blue. Compounds (in white) are scaled in size according to their number of connections within the network. Network depicted with Gephi version 0.9.2 using a ForceAtlas2 algorithm^88^ (settings: scaling=5.0, gravity=1.0, tolerance=1.0, approximation=1.2)**. B.** A simplified depiction of the number of reaction connections between (directed arrows) and within (circular arrows) each of the layers.

## Methods

### Generation of the translation and polymer biosynthesis reaction databases

The purposes of this study are (i) to describe translation as a chemosynthetic process, (ii) to integrate this process with a chemical reaction network describing metabolism; (iii) to map network-level patterns between object mass and object connectivity across different chemical domains; and (iv) to assess whether network-level patterns observed at the highest levels of the biotic hierarchy are present at its lowest levels of biochemistry. These purposes circumscribe the scope of the resulting network’s component description. Objects and reactions that describe the chemical components of the cellular translation and metabolic systems will be transcribed and included in the network. Those reactions that do not alter the explicit biochemical components of an *E. coli* cell (*i.e.,* gene regulatory interactions, sequence-specific motifs and their mutual interactions, isomerization reactions, *etc*.) may be otherwise important, but they fall outside the scope of this particular chemosynthetic study and will be excluded. To be sufficient for inclusion in the network, an object and its associated reactions must have been reported in peer-reviewed literature and be based on observable, experimental data; inferences based only on modeling alone are not sufficient.

To serve this purpose, the level of description for translation includes a chemical accounting of ribosome assembly and replication, as well as adjacent active site (A, P and E) interactions, since each of these interactions has downstream effects on the synthesis of biopolymers that enable autocatalytic replication of the entire system (**Figure 2B**). The chemical reaction network is intended to focus on the cellular chemosynthetic process, and does not explicitly include sequence-specific or site-specific (physiological or anatomical) information of the cell.

A list of *E. coli* translation components was obtained from the EcoCyc database^29^ (GO: 006412). **Supplementary File 1** summarizes the components of translation used in this network, apart from the ions and metabolites (*e.g.* GTPs) required to operate. The reactions for tRNA aminoacylation by tRNA synthetases, rRNA maturation and modifications, ribosomal protein modifications and maturations and peptidyl tRNA hydrolysis were obtained from the EcoCyc^29^ database and were also confirmed by literature synthesis (**Supplementary File 1**). The reactions for ribosomal subunit assembly, initiation, elongation and termination steps and stress response were built for *E. coli* (**Supplementary File 2**). The generation of translation reactions focused on five processes: a) elongation, which is largely conserved across all organisms; b) initiation, where some variations have been described between and within domains of life; c) termination; d) translation machinery biosynthesis, which are the assembly steps for replicating a ribosome; and e) the response to amino acid starvation. The reactions were manually annotated. To describe the elongation process in detail, the reaction model accounts for each of the three ribosome active sites. This includes a large number of reactions describing each of the distinct combinations of amino acids at each position. The metabolic and translation component masses were retrieved from the Kyoto Encyclopedia of Genes and Genomes (KEGG) Application Programming Interface (API). The TM layer reaction network file is attached as **Supplementary File 3**.

The polymer biosynthesis layer was constructed to link the metabolic and translation layers together by writing non-stoichiometric reactions that approximate the assembly of amino acids into the assembled enzymatic polypeptides. Each reaction within the metabolic layer associated with a specific enzyme resulted in a new reaction placed within the polymer biosynthesis layer network. For reactions that involved multiple protein components (*i.e.*, the formation of multimer heterocomplexes), the logical relationships of the protein components that enabled specific reactions were automatically read and parsed to generate protein-protein interaction maps within the molecular biosynthesis layer. The polymer biosynthesis layer is attached as **Supplementary File 6**.

The translation process is carried out as a repetition of steps for each of the proteins. Given that we are not considering sequence-specific processes or relationships, we must employ a placeholder “peptide” node that is used in chemosynthesis steps carried out by translation prior to completion of enzyme synthesis. It denotes any incomplete polypeptide sequence that is in the process of being assembled through the translation process. It does not possess any specific chemical function or identity that must be subsequently tracked in other portions of the chemical reaction network.

### Metabolic reaction database

The metabolic layer was obtained from Feist *et al.*^30^, who reconstructed *E. coli* metabolism (**Supplementary File 4**). We linked these reactions to their corresponding enzyme(s). We obtained chemical information about each of these enzymes through KEGG. The metabolic layer is attached as **Supplementary File 5.** We also considered metabolic models of alternative strains of *E. coli* obtained from BiGG^31^. The subtle differences between the strains do not lead to any significant difference in the degree distributions (see **Figure S1**), so we only consider one strain in this article.

### Chemical reaction network integration

All chemical reactions from each of the three layers (translation, polymer biosynthesis and metabolism) were uniformly processed into an aggregate network data file for graphing and analysis (**Figures 1 and 3A**). The aggregate network data file forms a bipartite graph with one class of nodes representing chemical compounds, and another class of nodes representing logical chemical relationships (*i.e.*, chemosynthesis reactions and biosynthesis assembly processes) that describe generating biomolecules from underlying substrates. The integrated network is composed of 1433 reactions for translation, 1455 reactions for polymer biosynthesis and 2085 reactions for cellular metabolism, each with varying proportions of intra- and internetwork connectivity. The complete network is provided in **Supplementary File 7.**

### Network visualization, processing, and analyses

We define the degree of a compound based on the number of different reactions involving it (whether as reactant, product, transporter or catalyst). We have built upon prior network analysis efforts^5^ to explicitly consider enzymes and macromolecular assemblies in addition to metabolites. The complementary cumulative distribution function (CCDF), a binned list of the number of compounds with a degree value in a given network, was analyzed using the power-law Python library^32^ to determine the likeliest statistical model that fits the distribution. We performed discrete and continuous fits using models of exponential and power-law distributions, and we compared the log-likelihood ratios among them to determine if a given degree distribution is likelier to be homogeneous or heterogeneous, respectively^33^.

Network visualizations with many pieces of information tend to become cluttered, which can obscure observational insights. Techniques that reduce a network’s information density without altering its fundamental relationships are useful for recovering these insights. We simplified network visualizations by grouping sets of reactions into combined nodes (*node-contracting*) according to their *module* labels. These labels were either provided by the original metabolic data without further modification^30^(*e.g.,* transport, pyruvate synthesis, *etc.*), or manually assigned by the authors for the translation network (*e.g.,* elongation, initiation, *etc.*). The term ‘module’ denotes a subset of nodes in a network that are densely connected to each other, while being sparsely connected or entirely disconnected from nodes in other modules^34, 35^. Compounds that participate only in reactions within a single module are grouped within combined nodes with their module labels, while those compounds that interact across different modules are conserved and graphed as individual nodes. The resulting contracted network is still a bipartite network with modules and reactions connected by compounds. We contracted a network formed by the translation and metabolism layers, and we perform an analysis to find out which nodes are more relevant in the traversal of this contracted network using the betweenness centrality metric. This metric consists of measuring the number of times that a node appears in the shortest path between two different nodes^36^:

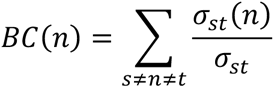

Where σ_st_ is the number of paths joining the nodes s and t, and σ_*s*t_(*n*) is the number of those paths that pass through the node n. Reaction directionality is accounted for when using this method. A different contraction approach is taken in Figure 3.B, where we collapse all the reactions belonging to the same layer, and we represent in the edges the number of compounds that participate within each layer or that connect the different layers.

The KEGG database provided the molecular structures and sequences required to compute the molecular weight of each of the chemical compounds (proteins, RNA, metabolites, *etc.*). In the case of complexes, we add the masses of their components but we disregard their stoichiometry of assembly due to a lack of experimental information.

Large networks are highly dimensional objects where similarity relationships are difficult to assess. Network embeddings allow the mathematical representation of network entities to ease their comparison^37^. The network embeddings are generated using the node2vec algorithm^38^ as implemented in PyTorch Geometric^39^. We use standard parameters to generate the embeddings. The resulting 128 dimensions of each of these embeddings were converted into 2D embeddings by using the t-distributed Stochastic Neighborhood Embedding (t-SNE) technique. Visualization of the integrated network and its individual layers was carried out in Gephi 0.9.2^40^. The code employed to perform this analysis together with the data are provided in the article GitHub repository https://github.com/kacarlab/Translation-Network.

## Results

Translation is a process described in textbooks in great detail, but there are still many unknowns about its function and modes of integration with other cellular systems. By reviewing the literature, we compiled the reactions of translation and generated a bipartite graph represented in **Figure 2B**. This TM network includes the core of translation processes (initiation, elongation and termination) together with the assembly and recycling of the ribosome, and the response to amino acid starvation. A feature of this network is the high level of redundancy of some of the reactions, given that translation is based on parallel processes involving different tRNAs.

All chemical reactions for translation and polymer biosynthesis were then integrated with that for metabolism. The chemical object names in each network were screened for consistent annotation, their molecular masses and object classifications were cross-referenced or inferred (*i.e.*, proteins, RNAs, codons, amino acids, energy molecules, and relevant ions). A full map of the integrated network, with node sizes weighted by node degree and reactions color-coded by network layer category is displayed in **Figure 3A**. The overall network contains more than 8000 nodes and 24000 links. When nodes are placed using a graph-layout algorithm (Force-Atlas 2 in this case), each layer clusters as a separated group of nodes that are mostly distinct from one another. A breakdown of the numbers of objects connected within and between the different layers is displayed through the use of numbered arrows in **Figure 3B**.

To understand how their components are connected within and outside each network, we applied Node2Vec non-supervised learning to generate multi-dimensional representations (embeddings) as vectors for each node in the network that reflect its neighborhood of connections. We then reduced these multi-dimensional vectors to two dimensions using a t-Distributed Stochastic Neighbor Embedding (t-SNE) dimensionality reduction technique. The resulting embeddings provide a representation of which nodes share a common neighborhood by placing them closer in a unitless 2D plane. The translation and the metabolic layers represent two different domains that can be easily distinguished (**Figure S2**), while the biosynthesis nodes appear scattered across these two different domains without any distinct clustering. Within the translation network, we find two large clusters of nodes and many smaller groups. Within these smaller groups there are nodes representing each of the amino acid processing cycles and the tRNA-synthetase process. In contrast, we do not find any cluster in the metabolic network containing distinct components of the TM.

Despite the large number of new connections mapped across the integrated network, the sparsity between translation and metabolic layers may be explained in part by the relatively few compounds joining them directly together (25 and 27 to and from each layer, **Figure 3B**), which is limited to the molecules providing peptide bond formation energy and substrate molecules for RNA polymers. It seems likely that the reactions that form connections across the metabolism and translation layers might have a central role keeping the overall network together. We contracted reactions and compounds involved in identified modules (*e.g.,* cofactor synthesis, elongation, *etc.*), and color-coded the resulting network using the betweenness centrality for each node of the contracted network in **Figure S3**. In this case, this metric represents how often a node is visited when traversing across the different layers of the integrated translation-metabolic network. The network depiction of these data shows a distinct separation between the translation and metabolism layers, with connections across the separation maintained through modules such as tRNA charging providing the raw materials for polypeptide elongation. Together with transport modules that provide the bulk of metabolic precursors for heterotrophic *E. coli* metabolism, tRNA charging is one of the most central modules of the cell.

An important attribute of large networks is their degree distribution: the probability of finding nodes with a given number of connections. In bipartite reaction networks, such as the integrated network assembled here, the degree of each compound represents the number of reactions in which it participates. Networks associated with complex systems usually show heterogeneous degree distributions, where a few nodes show a remarkable number of connections, while random networks are characterized by having more homogeneous degree distributions peaking around a mean number of connections^14^. We use the log-ratios of different statistical fits to determine the most likely degree distribution (exponential for homogeneous networks, power-law for heterogeneous networks). For objects that compose the metabolism and translation layers, their degree distributions follow statistically probable heavy-tailed distributions (**Figure 4B**, **Table 1**), which is not the case of the polymer biosynthesis layer degree distribution (p>0.05). The only object in the polymer biosynthesis network that is heavily connected to other compounds is a generic node ‘peptide’ (**Figure 3A**), which represents intermediate peptide sequences that for various reasons have not completed the termination step, and thus have no distinct chemosynthetic or functional impact on the network. The overall degree distribution of the full network follows a heavy-tailed degree distribution with an exponent around 2.0 (**Figure 4A**, **Table 1**).

**Figure 4.**
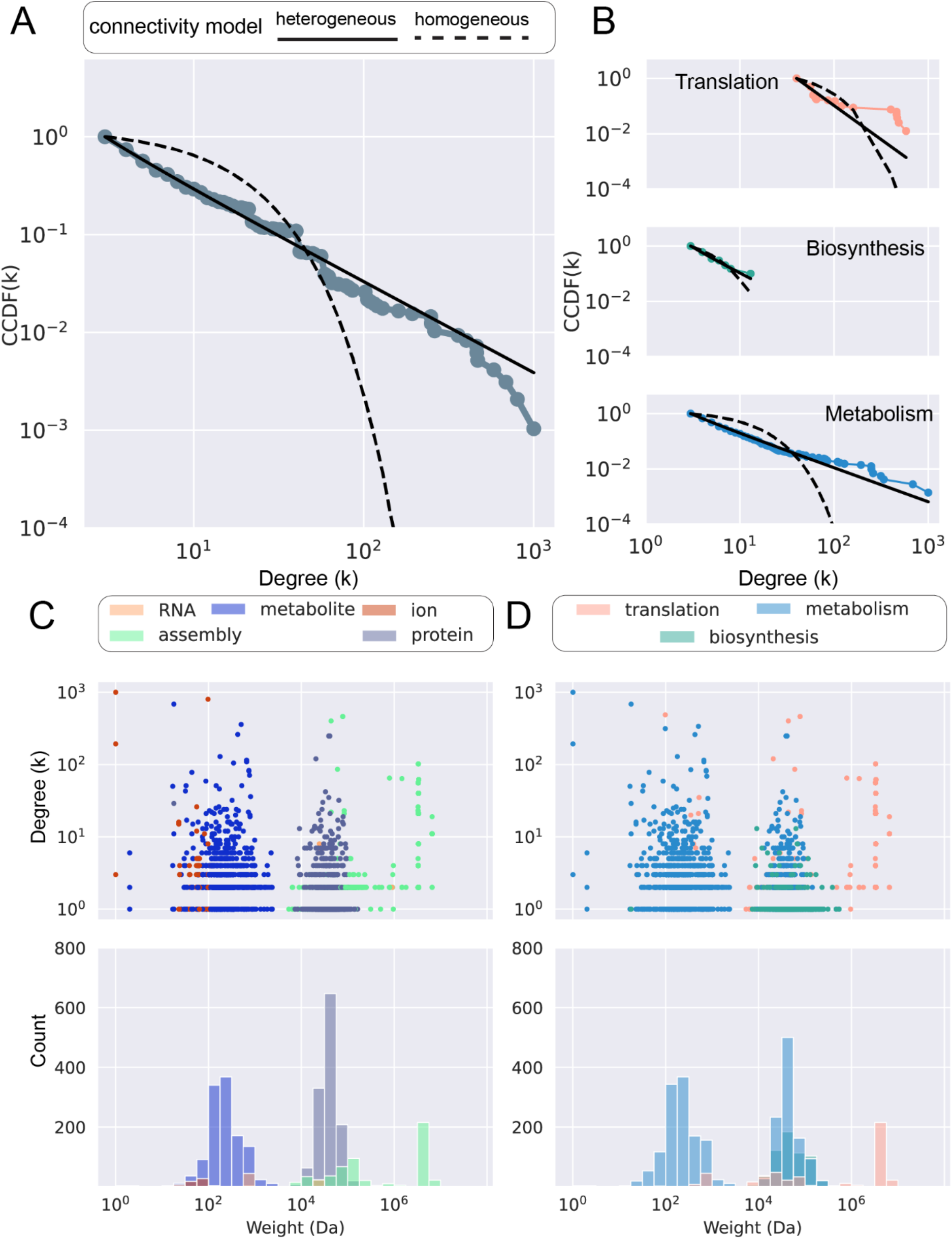
Statistical analysis of the degree and molecular weight data of the integrated chemosynthesis network and each of its layers. **A.** Complementary cumulative density function (CCDF) of node connection plotted versus bins of node connection degree (k) and distribution model comparisons for the complete chemosynthesis network; heterogeneous (exponential) model shown in dashed line, and heterogeneous (power law) model shown in solid line. **B.** CCDF plotted versus node connection degree (k) and distribution model comparisons for the Translation (top), Biosynthesis (middle) and Metabolism (bottom) layers; heterogeneous (exponential) model shown in dashed line, and heterogeneous (power law) model shown in solid line, as in Panel A. **C.** Raw scatter data of the connection degree (k) and molecular weight (Da) of all chemical objects in the network (upper panel) and same data depicted as a histogram of molecular weight bins (middle panel), with histogram bin contributions color coded by molecule type (bottom panel). **D.** Raw scatter data of the connection degree (k) and molecular weight (Da) of all chemical objects in the network (upper panel) and same data depicted as a histogram of molecular weight bins (middle panel), with histogram bin contributions color coded by layer (bottom panel).

**Table 1.** A tabulated comparison of power law and exponential distributions (log-likelihood ratios), significance estimates (p-value), minimal degree value at which we start fitting the distribution (k_min_), and best-fit exponent values (**γ**) for the complementary cumulative distribution functions of the translation, biosynthesis and metabolism networks. See Clauset *et al.*^33^. For a more detailed discussion of the fitting methods.

| Domain | Fit type |  |  |  |  |  |
| --- | --- | --- | --- | --- | --- | --- |
|  | Continuous |  |  | Discrete |  |  |
| | $\gamma$ | $k_{\min}$ | p-value | $\gamma$ | $k_{\min}$ | p-value |
| All layers | 2.118 | 3 | $2.4 \cdot 10^{-17}$ | 1.929 | 3 | $4.5 \cdot 10^{-13}$ |
| Biosynthesis | 3.470 | 3 | $9.9 \cdot 10^{-2}$ | 2.703 | 3 | $9.1 \cdot 10^{-1}$ |
| Metabolism | 2.290 | 6 | $9.0 \cdot 10^{-8}$ | 2.229 | 3 | $3.0 \cdot 10^{-7}$ |
| Translation | 3.526 | 40 | $3.2 \cdot 10^{-14}$ | 3.448 | 40 | $1.2 \cdot 10^{-13}$ |

While the degree of a compound might reflect its relevance in shaping the chemical network, its mass correlates with the amount of energy required for its synthesis; larger molecules require more energy to synthesize than smaller ones^41^. Our objective is to assess the relationships, if any, between the connection degrees and the masses of the compounds in the network. A multimodal distribution can be discerned in the panels of **Figure 4C** (categorized by chemical compound type) and **Figure 4D** (categorized by network layer). Mass distribution concentrations (modes) across all network layers occur at around 240, 40,000 and 2,000,000 Daltons (Da). The central regions of these modes contain both the highest concentrations of compounds and the most highly connected compounds from the different layers. There are bins in troughs between the modes without reported molecules, which indicates that the modal mass distribution may also be discontinuous. All modes contain some objects from the TM network; enzymatic polymers that enable metabolism are almost entirely confined to the middle mode; and metabolic compounds compose the two smallest modes. The structural composition of the chemical compounds (small molecules, polypeptides, and polynucleotides) is the likeliest determinant of the mode regardless of the functions they perform or the layer they are assigned to within this network (**Figures 4C and 4D**).

## Discussion

Bioinformatic databases are constructed and annotated in such a way as to enable the high-throughput reconstruction of many biosynthetic steps of metabolism and gene regulation^42^, and also to reconcile these steps with sequences found within an organism’s genome in an organized pipeline (gene → protein → function^43^). Prior network descriptions and analyses of translation have focused on its dynamic attributes afforded by its autocatalytic motifs^44^, genetic regulation of metabolic processes^45^, the links between TM function and organism-level cellular dynamics^46^ and a mapping of genotype to phenotype useful for interpreting a wealth of -omics cellular data^47^. The chemosynthetic function of the TM and the topological attributes of the overall network did not Figure Significantly into these prior efforts.

The integration of translation and metabolism reactions for *E. coli* describes a naturally occurring chemical network that is hierarchical and spans at least six orders of magnitude of compound mass. Hierarchy is most clearly resolved when degree per compound is plotted against compound mass (**Figure 4C and 4D, upper panels**). By inference, the energy costs associated with synthesizing and replicating the largest TM components outstrip those of enzymes and metabolites by at least 1-2 orders of magnitude. The energy cost per molecular component demonstrates the hierarchical importance of the TM in specifying chemical functions made possible by biopolymers within the cell.

As cellular operation requires a hierarchy of different mass entities, it also involves modules with different connectivity patterns. The embedding analysis depicted in **Figure S3** reveals that the metabolic and polymer biosynthesis reactions do not cluster apart from each other in any significant way, but the TM clusters in a distinct region of the graph. The metabolic and TM layers exhibit degree distributions that are likely heterogeneous, but the biopolymer layer does not. Taken together, this would indicate that protein biosynthesis is mostly shaped by the same structural constraints of the intracellular metabolism module and is not likely to exhibit independent (*i.e.*, intra-layer) attributes associated with complexity.

### Continuous, multimodal or discontinuous distributions

Based on prior observations of biological networks across the hierarchy of life, we articulated three distinct possibilities for the distribution of compounds in the integrated translation network (see **Figure S4**). Each of these possibilities has distinct, predicted implications that may be compared to the observed mass distribution. In a parallel to ‘trophic hierarchy’, one hypothesized driver of hierarchical connection in cellular biochemistry is the ability to distribute energy and maximize power in a dynamic way across independently functioning layers. For this possibility, each distinct molecular layer is self-organizing and both feeds upon and into the others. The predicted pattern is that each layer contains distinct connectivity peaks, and the overall mass distribution is continuous. In an ‘energy availability’ distribution, the hierarchical structures of polymer biosynthesis and translation are built directly atop, and extensions of, energy pathways enabled by metabolism. The predicted pattern is that though the translation and polymer biosynthesis layers of the network may contain ‘heavy tails’, the bulk of compounds in each layer all coincide with a peak generated by metabolic compounds. In a distribution shaped by ‘molecular distinction’, the association of compounds with a given biofunctional layer is less significant than the compounds’ chemical attributes in the cellular environment. The predicted pattern for this distribution is that multimodality or discontinuity caused by distinct molecular properties would lead to sorting at the network level. Each of these predicted distributions may carry basic implications about the primordial relationship(s) between emergent metabolic, enzymatic and translation functions, and about identifying measurable network attributes that fundamentally distinguish abiotic and biotic systems. We predicted that energy availability would be the dominant organizing driver of cellular biochemistry, given that energy transduction is indispensable to the overall function of translation.

The observed multimodal (and perhaps discontinuous) mass-degree distribution of the network seems most consistent with the ‘distinct molecules’ prediction. The network’s multimodal mass-degree distribution is the first such distribution reported for biological systems at the cellular level. The multimodal mass distribution of the integrated chemical reaction network is independently supported by chemical assays of living *E. coli* cells. Non-targeted metabolomic assays of small molecular mass compounds using mass spectrometry (TOF-MS) and different preparatory analytic weighting techniques consistently show a peak in metabolite compounds at approximately 150-350 Da, and a tapering of compounds approaching ∼2000 Da, even after accounting for decreased detector sensitivity with increasing mass^48, 49^. This approximately aligns with the centroid and distribution range of the smallest mass mode. For polypeptides, proteomic assays conducted across taxa as diverse as *E. coli, S. cerevisiae* and *H. sapiens* consistently show a broad correlation between translated genomic length and *in vivo* enzyme length^50^. For *E. coli*, this specifically includes a median peak length of about 209 amino acids, which corresponds to a peak in the mass distribution of polypeptides of approximately 28,000 Da if the average amino acid mass is assumed to be 136 Da (the average molar mass of all 20 essential amino acids). This approximately aligns with the centroid of the second mass mode. A whole-cell assay of all *E. coli* enzymes, regardless of composition and function, show that highly connected objects central to both metabolism and translation (*e.g.*, the ribosome, TCA cycle components, glycolysis and fermentation components, *etc.*) compose comparable protein mass fractions under a variety of growing conditions^51^. The ribosome itself is consistently composed of 2:1 mass fractions of RNA to protein^52^. In sum, independent raw cell molecular assays indicate sharply discretized compound mass modes that are attributable to the chemical properties of RNA polymers, proteinaceous polypeptides, and base metabolites, regardless of their cellular functions they carry out or the modules to which they may be assigned. Molecules that fall within the troughs between the modes are undoubtedly present in cells in some form, but would not seem to play essential roles in generating an *E. coli*’s chemosynthetic pathways, cellular physiology, or adaptive responses to stimuli. Multimodal distributions present a unique challenge to theoretical studies of network generation, and may require elaboration using multiplexes that can account for multimodality^53^.

### Causes and evolutionary implications of biochemical modularity

An assessment of the possible drivers of multimodality and discontinuity in the chemical reaction networks of bacteria may open new ways of characterizing the evolvability of cellular systems, and for distinguishing them from complex abiotic systems. Different network analysis techniques (**Figures 3A, S2 and S3**) and raw plots of mass distribution modes (**Figures 4C and 4D**) trace modularized systems of translation and metabolism that, by virtue of their chemical isolation from one another, are each able to independently adapt to conditions that the cell may encounter^48^. Modules can exhibit hierarchical properties and function as key units of adaptive evolution because organismal fitness depends on their performance^7, 54–57^.

The *E. coli* metabolic network is only able to exist via the functions afforded by polymeric peptides. Accounting for these polymeric peptides in the chemical reaction network introduces the first modal trough (**Figure 4C**) between the largest metabolites and the smallest polypeptides. This trough is also present in objects that compose the translation network that are found across this range of masses (**Figure 4D**). From a purely physical perspective, there are no *a priori* reasons for a discontinuity of mass size between these compounds. A discontinuity may be understandable, though, in biofunctional terms if resources for synthesis are limited^41^. The synthesis of an extremely large metabolite necessitates the precise placement of assorted atoms and molecular groups, connected with different bond types. If there is a single placement error, or inadequate feedstock of starting materials, the resulting compound is unlikely to be functional. An expenditure of chemical resources to produce increasingly larger (and increasingly error-prone) metabolites has adverse effects on the persistence of the overall cellular system. The synthesis of a linear oligopeptide of comparable mass, by contrast, can make use of repetitive interconnections of a much smaller number of object types (*i.e.*, amino acids), many of which also have similar chemical properties. A reduction of the number of building block types and of the kinds of bonds through which they are connected can increase production fidelity (and resultant functionality) of peptidic complexes compared to massive metabolites. The polypeptide assembly process cannot scale downward indefinitely, however, since there are natural limits to how small a chemically functional polypeptide may be. Oligopeptides must attain some minimal length to form structural folds, or to form a pocket in which specialized cofactor chemistry can occur that is isolated from the cellular environment^57^. The tradeoffs between synthesizing extremely large metabolites or extremely small polypeptides to ensure overall functionality can introduce a natural trough in mass distribution between these two chemical regimes that is attributable to both the constraints brought about by the chemical properties of their different building blocks and of the emergent functions they enable.

The trough between the second (polypeptide) and third (polynucleotide assemblies) modes appears to arise for entirely different reasons that are distinct to the TM. As with the first mode gap, there are no obvious physical or chemical reasons why there should not be an overlap between large polypeptide and small ribonucleotide polymer compounds, and the observed trough is counter to our predictions. The ribosome, however, has evolved to localize many of the TM’s protein-protein, protein-ribozyme, and ribosomal interactions on itself, thereby spatially isolating the functions of translation from the surrounding metabolic and physiological activities in the cell^55^. These functions are further isolated in time and space by regulatory mechanisms, such as via binding with the properly formed initiation factors^58–61^, by corresponding activated tRNA complexes^62^ and through spatial heterogeneity arising from complex physiological relationships with nucleoids^63^. The TM network description includes a rich array of intraribosomal subunit interactions, which we record as interactions involving an object with the mass of the ribosome. This creates a sharp, isolated peak in the distribution centered at the ribosome’s mass. The ready clustering of the TM in the network topology (**Figures 3A**) and embedded vector and dimensionality reduction analysis (**Figure S2**) depict and corroborate the biochemical facets of translation’s modularity, as do *in vitro* experimental systems that employ cell-free protein synthesis^64, 65^.

There are numerous examples of systems that may be characterized as complex self-organizing, adaptive phenomena, but which lack the evolvability, emergent novelty and agency of living systems. Far-from-equilibrium physical systems such as galactic, stellar, and planetary structures^66^, earthquakes^67^, solar flares ^68^ and sandpile or avalanche dynamics^69^ all exhibit continuous scaling across many orders of magnitude, but few or no reported discontinuities. Discrete size classes observed in biology, on the other hand, are evidence of multiple dynamic regimes^70^ in which signaling, and the perturbative effects of novelty can propagate and be controlled in directions both up and down scales of a hierarchy^71^.

A multimodal biochemical distribution may be interpreted as indicating an evolved response of modularization to significant constraints placed on a complex adaptive chemical reaction network^34^. In biochemical systems, energy flows through the entire hierarchy mostly in the direction from metabolites up to the translation components, but modal gaps and troughs arise due to a complicated reconciliation process between the basic building blocks that compose large objects in the system, energy availability for synthesis, and optimization of functional traits that large objects can exhibit^72^. In conjunction with a heavy-tailed degree distribution, a multimodal mass distribution may be one of a select suite of indicators diagnostic of biological dynamics that can be observed across the entirety of life’s various spatiotemporal scales.

### Implications for prebiotic chemistry

Mapping the full extent of chemosynthetic relationships between translation and metabolism (**Figures 3A** and **3B**) graphically captures long-recognized aspects of the hierarchical relationship between these two cellular functions. Translation of a transcripted mRNA directs the specificity of polymer-catalyzed metabolic chemistry, and this chemistry in turn supplies the required energy and materials for ribosomal replication and operation. At the same time, though, the existence of multiple distinct mass discontinuities that correlate more with differences in chemical composition (**Figure 4D**) than with cellular function (**Figure 4C**) opens new possibilities about the underlying dynamic circumstances in which translation arose. Chemical modes and discontinuities may be so fundamental as to have been contemporaneous with (or have preceded) the origins of cellular life. As an extension, it is possible that some information-processing attributes of the TM module are primordially rooted in complex feedbacks between ribonucleotides and metabolites, intra-network dynamics of the TM, or physicochemical reinforcement of modularization (such as spatiotemporal isolation) afforded by discrete compound class modes, as much as from the emergence of the ribosome’s large and small subunit mechanical activities.

Generating hierarchically connected networks across so many orders of magnitude, with modules that each contain at least one autocatalytic cycle^44, 73–75^ presents a significant challenge to the field of prebiotic chemistry^76–78^. If the biochemical modality and modularity observed in this network are genuinely universal attributes of life, they may arise due to general physical phenomena such as percolation^79^. The observed coupling between translation, protein biosynthesis and metabolic networks presents a new model system in which the network phenomenon of percolation may be studied in a chemical context^80^ as an exemplary emergent system in which multiple networks with distinct topologies coexist and are connected via common elements^81^ and discontinuities stem from mechanisms that lead to network (dis)assortativity^82, 83^. Theoretical analyses of such conditions show a possible connection between the number of distinct modes and the possibility of multiple discontinuous phase transitions enabled by multiplex networks^84^. Phase transitions have long been recognized as potentially important behavioral features that are correlated with complex adaptive systems^85^.

The relationships between complex adaptive systems and information processing attributes afforded by network topology are directly relevant to developing new theories of the origin of translation. The decoding process that links nucleotide sequence identity to peptide sequence functionality is highly optimized for error tolerance through a property known as code degeneracy, which can be described in informatic and computational terms without explicit reference to the underlying biochemistry. It is unclear how an inherently informatic, network-level property such as code degeneracy would arise on the basis of tracing the individual origins of the ribosome’s chemical components and interaction partners in isolation from this wider network^86^. Recent discoveries have shown that ‘biological’ attributes such as self-reproduction, adaptation and evolvability (which together characterize much of biological agency) are intrinsically linked to one another through coupling of Hopf and Turing instabilities within a small network of reactions with a cubic autocatalytic motif^87^. These attributes are observable even in the absence of the intricacies of macromolecular mechanisms or a diversified biochemical metabolic network. Further studies of coupled metabolic and translation chemical reaction network properties may help to develop new hypotheses about the origin of translation as an information-processing modular network, rather than about how translation can arise due to the chemical relationships between amino acids and nucleotides or through the specific biochemical activities of TM components. Specifically, one may test whether the modularization of information processing centered on the ribosome in modern cells can be decentralized across a primordial TM network.

## Conclusions

Life exists, in part, by cohesively channeling energy from abundant, small-mass objects into the synthesis of a specific array of large-mass objects. These larger objects afford functions and activities that smaller objects, for various reasons, cannot perform. The inherent tension between utilizing available energy in an efficient way and ensuring diverse functions and activities will be reliably carried out can lead to multimodal mass distributions. Multimodality in this sense may be diagnostically biological, as an evolved trait that is exhibited by adaptive, self-organizing and self-assembling populations that can exhibit and propagate novel responses to stimuli and which are subject to natural selection. It is a property that has been observed in many different ecological systems. Our study of a combined network of metabolism and translation shows that mass multimodality can arise in biochemistry, with distinct drivers for different troughs between modes, even before physiological or gene regulatory mechanisms of cellular operation are considered. This extends multimodality down to the lowest levels at which biological population dynamics can exist: the chemistry within individual bacterial cells. Translation stands apart in the cell as a functional module that is as much informatic as it is chemosynthetic. Its modularity is achieved in part by concentrating many enzyme and ribozyme component interactions on the ribosomal complex, which has the effect of chemically isolating the translation function from much of the chemical activity going on elsewhere in the cell. Looking forward, it may be possible to use an integrated metabolism and translation network to study the emergence of the translation function as a network-level module in a way that is abstracted from the mechanistic biochemical activities carried out by the ribosome and its interaction partners.

## Acknowledgements

This work is supported by the Wisconsin Alumni Research Foundation, the John Templeton Foundation Grant #61926 and the Universidad Politécnica de Madrid (UPM) Margarita Salas Fellowship, founded by the “Unión Europea-NextGeneration EU“ (code UP2021-035).

## Code availability

The code necessary to reproduce this study can be found in the article Github repository: https://github.com/kacarlab/Translation-Network

## Data availability

The data of this study can be found in the article Github repository: https://github.com/kacarlab/Translation-Network

## Author contributions

B.C.Z., E.F., Z.R.A. and B.K. conceptualized and designed the study. B.C.Z. and E. F. performed the experiment and data analysis with input from Z.R.A. and B.K. B.C.Z. and B.K. wrote the paper with input from E.F. and Z.R.A. B.K. supervised the project. All authors read and approved the manuscript.

## Supplementary Figures

**Figure S1.**
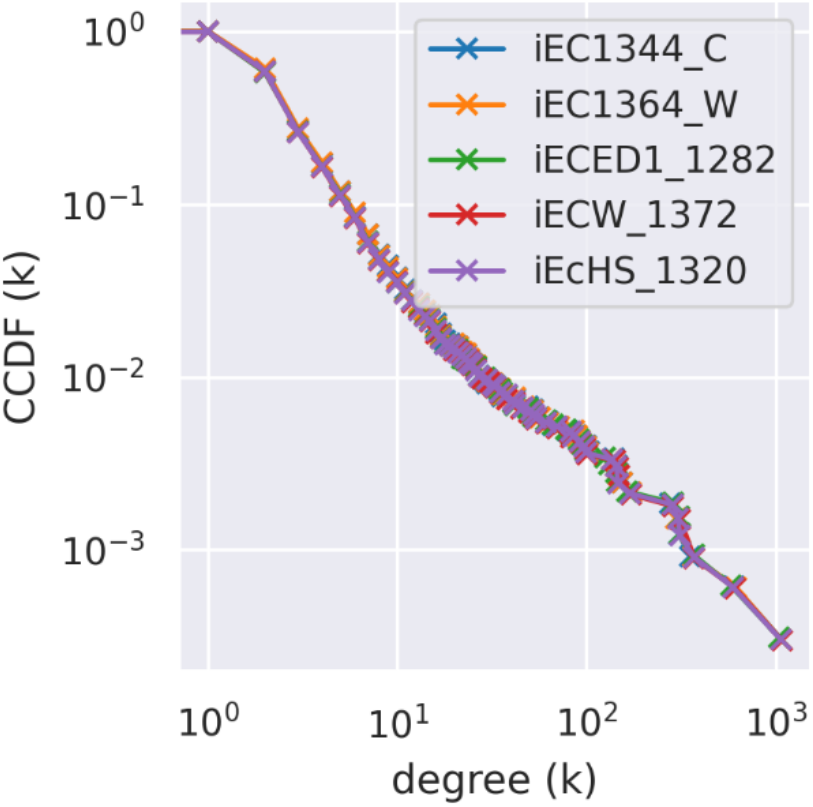
Degree complementary cumulative distribution function (CCDF) of five different strains.

**Figure S2.**
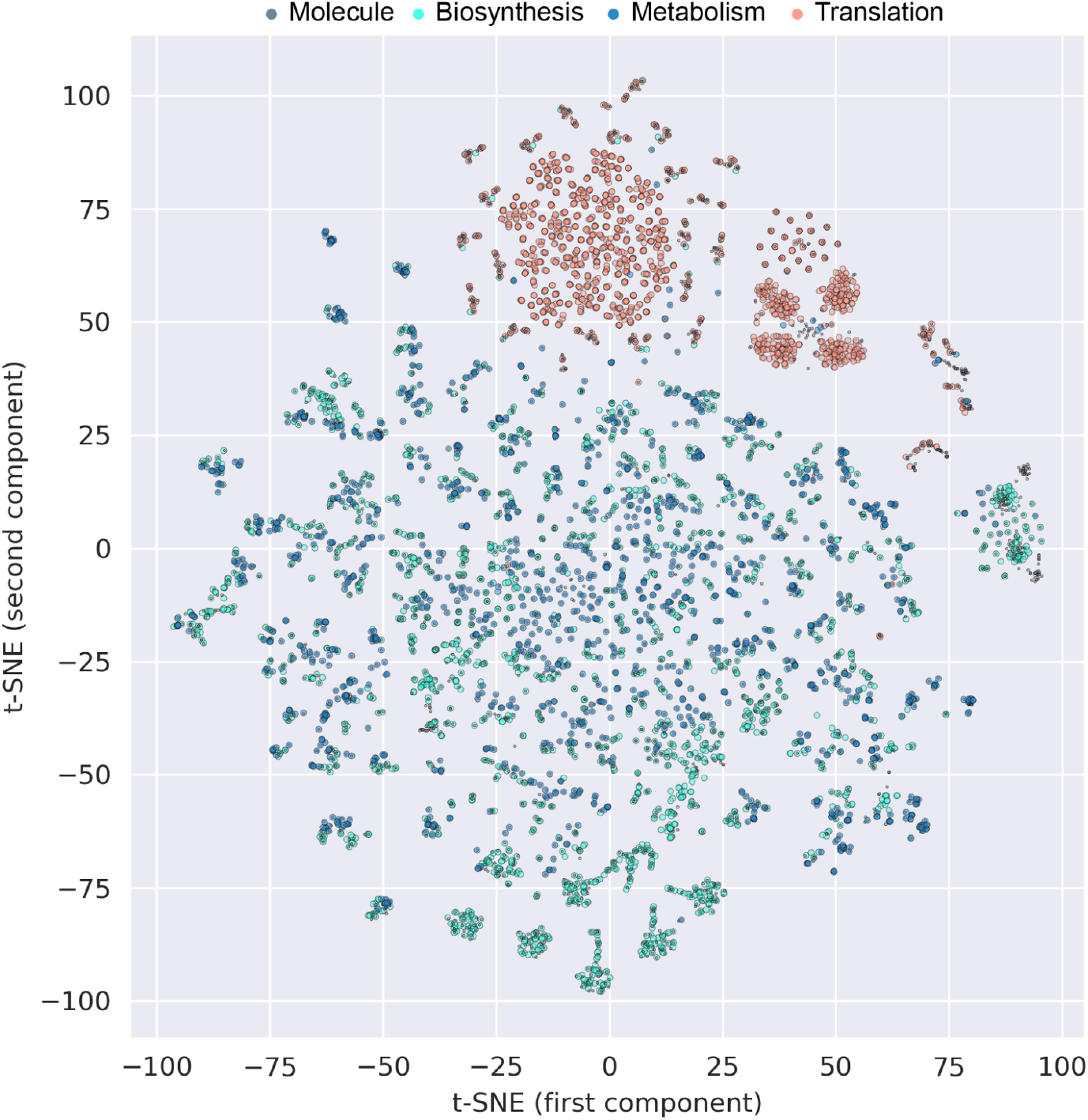
Scatter plot of the t-distributed stochastic neighbor embedding (t-SNE) dimensional reduction of node2vec embeddings of reactions from the complete integrated network. Reactions within the different layers are color coded with all chemical compounds depicted in gray and layer reactions from Translation in orange, Biosynthesis in green and Metabolism in blue. Reactions from the translation and metabolism layers group distinctly from one another with little overlap. Reactions from the biosynthesis layer do not group together in any distinct way, but they broadly plot in the same areas that contain metabolic reactions.

**Figure S3.**
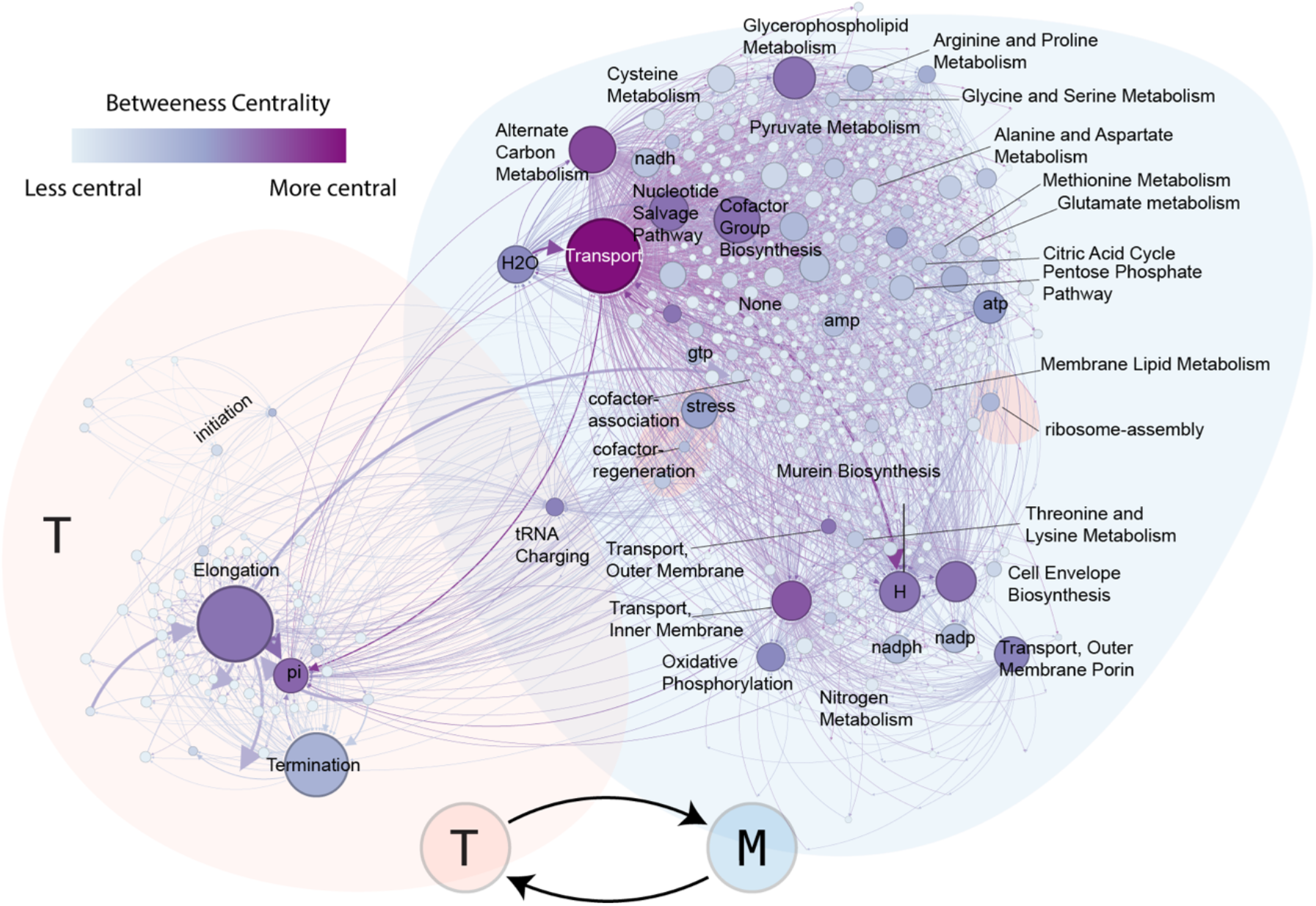
Simplified view of the integrated network depicting the connections between translation and metabolism modules. A module is a collection of reactions and compounds that only connect to other reactions and compounds within the same assignment category. Compounds that occur in more than one module are conserved (*i.e.*, not reduced to a module). Module categories for translation were assigned during layer construction using labels as shown in Figure 2B, while those for metabolism were taken without modification or alteration from the source document from Feist et al ^30^. Modules are coded from white (less central) to purple (more central) using the betweenness centrality metric; the more visited a node is in a path between two random nodes, the higher its betweenness-centrality.

**Figure S4.**
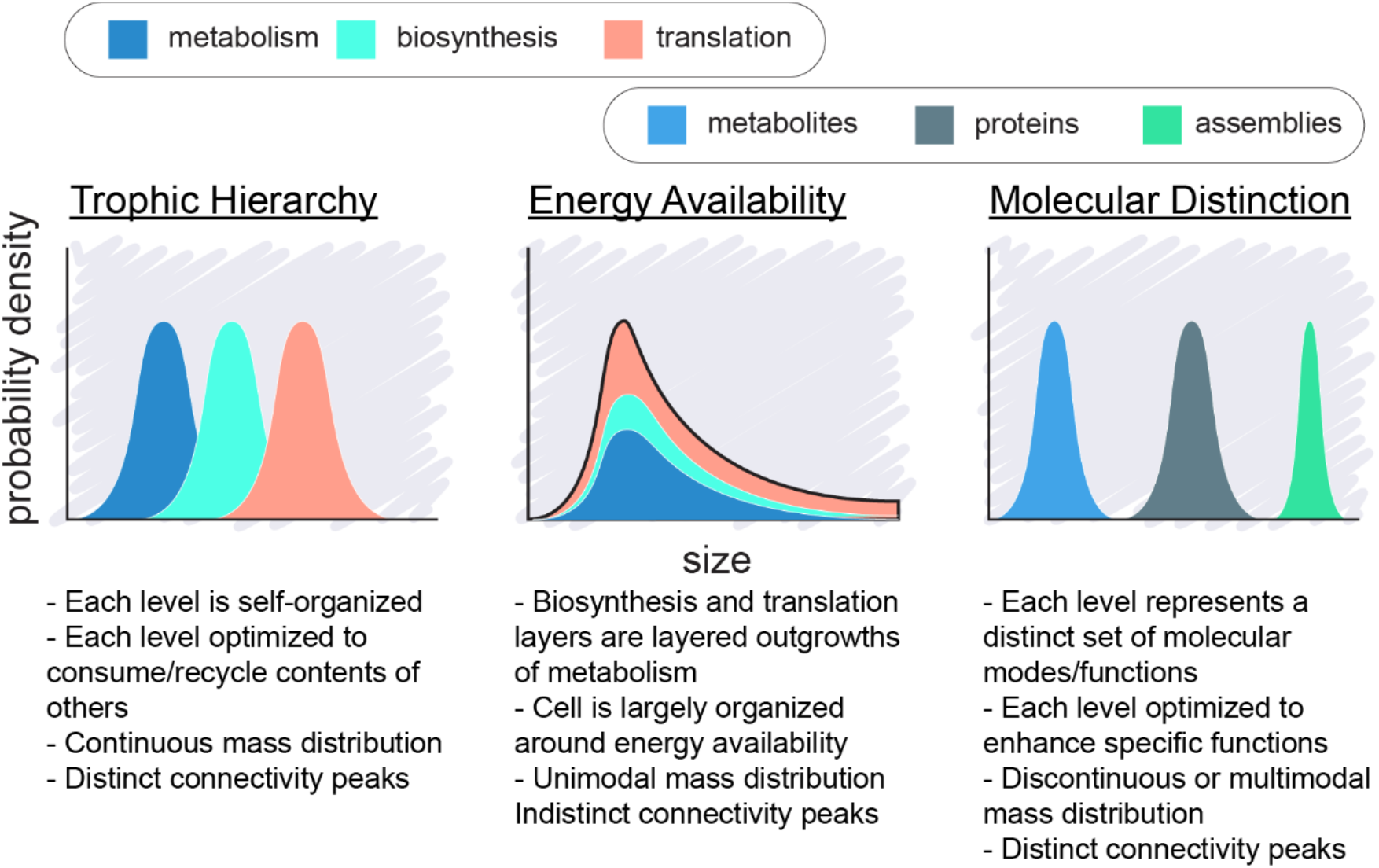
Depiction of three different expected patterns of compound size distributions (top panels) and their respective associated characteristics (bottom panels).

## References

1. Oberhardt, M. A., Palsson, B. Ø. & Papin, J. A. Applications of genome-scale metabolic reconstructions. Mol Syst Biol 5, 320–320 (2009).

2. Lewis, M. A Tale of Two Repressors. J Mol Biol 409, 14–27 (2011).

3. Chubukov, V., Gerosa, L., Kochanowski, K. & Sauer, U. Coordination of microbial metabolism. Nat Rev Microbiol 12, 327–340 (2014).

4. Chubukov, V., Zuleta, I. A. & Li, H. Regulatory architecture determines optimal regulation of gene expression in metabolic pathways. Proc National Acad Sci 109, 5127–5132 (2012).

5. Jeong, H., Tombor, B., Albert, R., Oltvai, Z. N. & Barabási, A.-L. The large-scale organization of metabolic networks. Nature 407, 651–654 (2000).

6. Barabási, A.-L., Albert, R. & Jeong, H. Mean-field theory for scale-free random networks. Phys Statistical Mech Appl 272, 173–187 (1999).

7. Ravasz, E., Somera, A. L., Mongru, D. A., Oltvai, Z. N. & Barabási, A.-L. Hierarchical Organization of Modularity in Metabolic Networks. Science 297, 1551–1555 (2002).

8. Newman, M. E. J. Analysis of weighted networks. Phys Rev E 70, 056131 (2004).

9. Ma, W., Trusina, A., El-Samad, H., Lim, W. A. & Tang, C. Defining Network Topologies that Can Achieve Biochemical Adaptation. Cell 138, 760–773 (2009).

10. Nghe, P. et al. Prebiotic network evolution: six key parameters. Mol Biosyst 11, 3206–3217 (2015).

11. Wagner, A. & Fell, D. A. The small world inside large metabolic networks. Proc Royal Soc Lond Ser B Biological Sci 268, 1803–1810 (2001).

12. Newman, M. E. J. The Structure and Function of Complex Networks. Siam Rev 45, 167–256 (2003).

13. Shenhav, B., Solomon, A., Lancet, D. & Kafri, R. Transactions on Computational Systems Biology I. in 14–27 (2005). doi:10.1007/978-3-540-32126-2_2.

14. Albert, R. & Barabási, A.-L. Statistical mechanics of complex networks. Rev. Mod. Phys. 74, 47–97 (2002).

15. Holling, C. S. Cross-Scale Morphology, Geometry, and Dynamics of Ecosystems. Ecol Monogr 62, 447–502 (1992).

16. Fischer, J., Lindenmayer, D. B. & Montague-Drake, R. The role of landscape texture in conservation biogeography: a case study on birds in south-eastern Australia. Divers Distrib 14, 38–46 (2008).

17. Havlicek, T. D. & Carpenter, S. R. Pelagic species size distributions in lakes: Are they discontinuous? Limnol Oceanogr 46, 1021–1033 (2001).

18. Lambert, T. D., Malcolm, J. R. & Zimmerman, B. L. Amazonian small mammal abundances in relation to habitat structure and resource abundance. J Mammal 87, 766–776 (2006).

19. Stow, C., Allen, C. & Garmestani, A. Evaluating Discontinuities in Complex Systems: Toward Quantitative Measures of Resilience. Ecol Soc 12, (2007).

20. Nash, K. L. et al. Discontinuities, cross-scale patterns, and the organization of ecosystems. Ecology 95, 654–667 (2014).

21. Novozhilov, A. S., Wolf, Y. I. & Koonin, E. V. Evolution of the genetic code: partial optimization of a random code for robustness to translation error in a rugged fitness landscape. Biol Direct 2, 24 (2007).

22. Koonin, E. V. & Novozhilov, A. S. Origin and Evolution of the Universal Genetic Code. Annu Rev Genet 51, 1–18 (2016).

23. Zaher, H. S. & Green, R. Quality control by the ribosome following peptide bond formation. Nature 457, 161–166 (2009).

24. Steinchen, W., Zegarra, V. & Bange, G. (p)ppGpp: Magic Modulators of Bacterial Physiology and Metabolism. Front Microbiol 11, 2072 (2020).

25. Prossliner, T., Gerdes, K., Sørensen, M. A. & Winther, K. S. Hibernation factors directly block ribonucleases from entering the ribosome in response to starvation. Nucleic Acids Res 49, gkab017- (2021).

26. Starosta, A. L., Lassak, J., Jung, K. & Wilson, D. N. The bacterial translation stress response. Fems Microbiol Rev 38, 1172–1201 (2014).

27. Irving, S. E., Choudhury, N. R. & Corrigan, R. M. The stringent response and physiological roles of (pp)pGpp in bacteria. Nat Rev Microbiol 19, 256–271 (2021).

28. Fuente, I. M. D. la, Cortes, J. M., Pelta, D. A. & Veguillas, J. Attractor Metabolic Networks. Plos One 8, e58284 (2013).

29. Keseler, I. M. et al. The EcoCyc Database in 2021. Front Microbiol 12, 711077 (2021).

30. Feist, A. M. et al. A genome-scale metabolic reconstruction for Escherichia coli K-12 MG1655 that accounts for 1260 ORFs and thermodynamic information. Mol Syst Biol 3, 121 (2007).

31. King, Z. A. et al. BiGG Models: A platform for integrating, standardizing and sharing genome-scale models. Nucleic Acids Res. 44, D515–D522 (2016).

32. Alstott, J., Bullmore, E. & Plenz, D. powerlaw: A Python Package for Analysis of Heavy-Tailed Distributions. Plos One 9, e85777 (2014).

33. Clauset, A., Shalizi, C. R. & Newman, M. E. J. Power-Law Distributions in Empirical Data. SIAM Rev. 51, 661–703 (2009).

34. Hintze, A. & Adami, C. Evolution of Complex Modular Biological Networks. Plos Comput Biol 4, e23 (2008).

35. Srivastava, A., Kumar, S. & Ramaswamy, R. Two-layer modular analysis of gene and protein networks in breast cancer. Bmc Syst Biol 8, 81 (2014).

36. Girvan, M. & Newman, M. E. J. Community structure in social and biological networks. Proc. Natl. Acad. Sci. 99, 7821–7826 (2002).

37. Nelson, W. et al. To Embed or Not: Network Embedding as a Paradigm in Computational Biology. Frontiers Genetics 10, 381 (2019).

38. Grover, A. & Leskovec, J. node2vec: Scalable Feature Learning for Networks. Arxiv (2016).

39. Fey, M. & Lenssen, J. E. Fast Graph Representation Learning with PyTorch Geometric. Arxiv (2019) doi:10.48550/arxiv.1903.02428.

40. Bastian, M., Heymann, S. & Jacomy, M. Gephi: An Open Source Software for Exploring and Manipulating Networks. Proc Int Aaai Conf Web Soc Media 3, 361–362 (2009).

41. Hu, X.-P., Dourado, H., Schubert, P. & Lercher, M. J. The protein translation machinery is expressed for maximal efficiency in Escherichia coli. Nat Commun 11, 5260 (2020).

42. Bar-Joseph, Z. et al. Computational discovery of gene modules and regulatory networks. Nat Biotechnol 21, 1337–1342 (2003).

43. Becker, S. A. & Palsson, B. O. Context-Specific Metabolic Networks Are Consistent with Experiments. Plos Comput Biol 4, e1000082 (2008).

44. Roy, A., Goberman, D. & Pugatch, R. A unifying autocatalytic network-based framework for bacterial growth laws. P Natl Acad Sci Usa 118, e2107829118 (2021).

45. Grimbs, A., Klosik, D. F., Bornholdt, S. & Hütt, M.-T. A system-wide network reconstruction of gene regulation and metabolism in Escherichia coli. Plos Comput Biol 15, e1006962 (2019).

46. Thiele, I. et al. Multiscale Modeling of Metabolism and Macromolecular Synthesis in E. coli and Its Application to the Evolution of Codon Usage. Plos One 7, e45635 (2012).

47. Thiele, I., Jamshidi, N., Fleming, R. M. T. & Palsson, B. Ø. Genome-Scale Reconstruction of Escherichia coli’s Transcriptional and Translational Machinery: A Knowledge Base, Its Mathematical Formulation, and Its Functional Characterization. Plos Comput Biol 5, e1000312 (2009).

48. Fuhrer, T., Heer, D., Begemann, B. & Zamboni, N. High-Throughput, Accurate Mass Metabolome Profiling of Cellular Extracts by Flow Injection–Time-of-Flight Mass Spectrometry. Anal Chem 83, 7074–7080 (2011).

49. Dwivedi, P. et al. Metabolic profiling of Escherichia coli by ion mobility-mass spectrometry with MALDI ion source. J. Mass Spectrom. 45, 1383–1393 (2010).

50. Geiger, T., Wehner, A., Schaab, C., Cox, J. & Mann, M. Comparative Proteomic Analysis of Eleven Common Cell Lines Reveals Ubiquitous but Varying Expression of Most Proteins*. Mol Cell Proteomics 11, M111.014050 (2012).

51. Mori, M. et al. From coarse to fine: the absolute Escherichia coli proteome under diverse growth conditions. Mol Syst Biol 17, e9536 (2021).

52. Kostinski, S. & Reuveni, S. Ribosome Composition Maximizes Cellular Growth Rates in E. coli. Phys Rev Lett 125, 028103 (2020).

53. Bianconi, G. Statistical mechanics of multiplex networks: Entropy and overlap. Phys Rev E 87, 062806 (2013).

54. Venkataram, S., Monasky, R., Sikaroodi, S. H., Kryazhimskiy, S. & Kacar, B. Evolutionary stalling and a limit on the power of natural selection to improve a cellular module. Proc National Acad Sci 117, 18582–18590 (2020).

55. Hartwell, L. H., Hopfield, J. J., Leibler, S. & Murray, A. W. From molecular to modular cell biology. Nature 402, C47–C52 (1999).

56. Qi, Y. & Ge, H. Modularity and Dynamics of Cellular Networks. Plos Comput Biol 2, e174 (2006).

57. Goldman, A. D. & Kacar, B. Cofactors are Remnants of Life’s Origin and Early Evolution. J Mol Evol 89, 127–133 (2021).

58. Simonetti, A. et al. A structural view of translation initiation in bacteria. Cell Mol Life Sci 66, 423 (2008).

59. Sharma, I. M. & Woodson, S. A. RbfA and IF3 couple ribosome biogenesis and translation initiation to increase stress tolerance. Nucleic Acids Res 48, 359–372 (2019).

60. Milón, P. & Rodnina, M. V. Kinetic control of translation initiation in bacteria. Crit Rev Biochem Mol 47, 334–348 (2012).

61. Benelli, D. & Londei, P. Begin at the beginning: evolution of translational initiation. Res Microbiol 160, 493–501 (2009).

62. Schuette, J. et al. GTPase activation of elongation factor EF-Tu by the ribosome during decoding. Embo J 28, 755–765 (2009).

63. Bakshi, S., Choi, H. & Weisshaar, J. C. The spatial biology of transcription and translation in rapidly growing Escherichia coli. Front Microbiol 6, 636 (2015).

64. Hartman, M. C. T., Josephson, K., Lin, C.-W. & Szostak, J. W. An Expanded Set of Amino Acid Analogs for the Ribosomal Translation of Unnatural Peptides. Plos One 2, e972 (2007).

65. Niederholtmeyer, H., Stepanova, V. & Maerkl, S. J. Implementation of cell-free biological networks at steady state. Proc National Acad Sci 110, 15985–15990 (2013).

66. Pérez-Mercader, J. Astrobiology, The Quest for the Conditions of Life. 337–360 (2002) doi:10.1007/978-3-642-59381-9_22.

67. Gutenberg, B. & Richter, C. F. Frequency of earthquakes in California*. B Seismol Soc Am 34, 185–188 (1944).

68. Krucker, S. & Benz, A. O. Energy Distribution of Heating Processes in the Quiet Solar Corona. Astrophysical J Lett 501, L213–L216 (1998).

69. Bak, P., Tang, C. & Wiesenfeld, K. Self-organized criticality. Phys Rev A 38, 364–374 (1988).

70. Garmestani, A., Allen, C. & Gunderson, L. Panarchy: Discontinuities Reveal Similarities in the Dynamic System Structure of Ecological and Social Systems. Ecol Soc 14, (2009).

71. Holling, C. S. & Gunderson, L. H. Resilience and adaptive cycles. *In:* Panarchy: Understanding Transformations in Human and Natural Systems, 25-62 (2002).

72. Shekhtman, L. M., Shai, S. & Havlin, S. Resilience of networks formed of interdependent modular networks. New J Phys 17, 123007 (2015).

73. Zubarev, D. Y., Rappoport, D. & Aspuru-Guzik, A. Uncertainty of Prebiotic Scenarios: The Case of the Non-Enzymatic Reverse Tricarboxylic Acid Cycle. Sci Rep-uk 5, 8009 (2015).

74. Braakman, R. & Smith, E. The compositional and evolutionary logic of metabolism. Phys Biol 10, 011001 (2013).

75. Preiner, M. et al. Catalysts, autocatalysis and the origin of metabolism. Interface Focus 9, 20190072 (2019).

76. Smith, J. I., Steel, M. & Hordijk, W. Autocatalytic sets in a partitioned biochemical network. J Syst Chem 5, 2 (2014).

77. Orgel, L. In the beginning. Nature 439, 915–915 (2006).

78. Shapiro, R. Small Molecule Interactions were Central to the Origin of Life. Q Rev Biology 81, 105–126 (2006).

79. Sornette, D. Critical Phenomena in Natural Sciences, Chaos, Fractals, Self Organization and Disorder: Concepts and Tools. Springer Series Syne 239–256 (2000) doi:10.1007/978-3-662-04174-1_12.

80. Schwartz, N., Cohen, R., ben-Avraham, D., Barabási, A.-L. & Havlin, S. Percolation in directed scale-free networks. Phys Rev E 66, 015104 (2002).

81. Leicht, E. A. & D’Souza, R. M. Percolation on interacting networks. Arxiv (2009) doi:10.48550/arxiv.0907.0894.

82. Newman, M. E. J. & Girvan, M. Finding and evaluating community structure in networks. Phys Rev E 69, 026113 (2004).

83. Newman, M. E. J. Mixing patterns in networks. Phys Rev E 67, 026126 (2003).

84. Kryven, I. & Bianconi, G. Enhancing the robustness of a multiplex network leads to multiple discontinuous percolation transitions. Phys Rev E 100, 020301 (2019).

85. Paperin, G., Green, D. G. & Sadedin, S. Dual-phase evolution in complex adaptive systems. J Roy Soc Interface 8, 609–629 (2011).

86. Gonzalez, D. L., Giannerini, S. & Rosa, R. On the origin of degeneracy in the genetic code. Interface Focus 9, 20190038 (2019).

87. Muñuzuri, A. P. & Pérez-Mercader, J. Unified representation of Life’s basic properties by a 3-species Stochastic Cubic Autocatalytic Reaction-Diffusion system of equations. Phys Life Rev 41, 64–83 (2022).

88. Jacomy, M., Venturini, T., Heymann, S. & Bastian, M. ForceAtlas2, a Continuous Graph Layout Algorithm for Handy Network Visualization Designed for the Gephi Software. Plos One 9, e98679 (2014).

